# Heritable epigenetic variation facilitates long-term maintenance of epigenetic and genetic variation

**DOI:** 10.1101/2022.12.19.521105

**Authors:** Amy K. Webster, Patrick C. Phillips

**Affiliations:** Institute of Ecology and Evolution, University of Oregon, Eugene, OR 97403

## Abstract

The maintenance of genetic and phenotypic variation has long been one of the fundamental questions in population and quantitative genetics. A variety of factors have been implicated to explain the maintenance of genetic variation in some contexts (*e.g*. balancing selection), but the potential role of epigenetic regulation to influence population dynamics has been understudied. It is well recognized that epigenetic regulation, including histone methylation, small RNA expression, and DNA methylation, helps to define differences between cell types and facilitate phenotypic plasticity. In recent years, empirical studies have shown the potential for epigenetic regulation to also be heritable for at least a few generations without selection, raising the possibility that differences in epigenetic regulation can act alongside genetic variation to shape evolutionary trajectories. Like genetic mutation, heritable differences in epigenetic regulation can arise spontaneously; these are termed ‘epimutations’. Epimutations differ from genetic mutations in two key ways – they occur at a higher rate and the loci at which they occur often revert back to their original state within a few generations. Here, we present an extension of the standard population-genetic model with selection to incorporate epigenetic variation arising via epimutation. Our model assumes a diploid, sexually reproducing population with random mating. In addition to spontaneous genetic mutation, we included parameters for spontaneous epimutation and back-epimutation, allowing for four potential epialleles at a single locus (two genetic alleles, each with two epigenetic states), each of which affect fitness. We then analyzed the conditions under which stable epialleles were maintained. Our results show that highly reversible epialleles can be maintained in long-term equilibrium under neutral conditions in a manner that depends on the epimutation and back-epimutation rates, which we term epimutation-back-epimutation equilibrium. On the other hand, epialleles that compensate for deleterious mutations cause deviations from the expectations of mutation-selection balance by a simple factor that depends on the epimutation and back-epimutation rate. We also numerically analyze several sets of fitness parameters for which large deviations from mutation-selection balance occur. Together, these results demonstrate that transient epigenetic regulation may be an important factor in the maintenance of both epigenetic and genetic variation in populations.

## INTRODUCTION

Epigenetic regulation is integral to the genotype-phenotype map. Indeed, the concept of the ‘epigenotype’ was originally defined to include anything ‘between genotype and phenotype’ (Waddington C 1942). However, modern usage is more limited to include either 1) specific heritable (mitotically or meiotically) modifications to DNA or RNA and/or 2) heritable changes in phenotype that are not dependent on an underlying nucleotide sequence. Studies employing the first usage commonly profile a population of cells or organisms for a specific epigenetic modification (Mehrmohamadi et al. 2021), under the assumption that epigenetic marks represent mitotically-heritable states and may impact gene expression. In contrast, the latter, more classical, definition does not require a specific type of modification but does require that a phenotypic change is inherited independently of genetic variation, with epigenetic modifications being likely candidates. This mode of transmission been used to denote transgenerational epigenetic inheritance in which an environmentally-induced phenotype is inherited across multiple generations without the initial trigger, even when the epigenetic mechanism is not known (Jobson et al. 2015; Webster et al. 2018). Typically, these examples of transgenerational epigenetic inheritance require that a phenotypic change is inherited three or more generations to ensure that inheritance is not driven by a maternal effect (*e.g*., yolk provisioning) (Burton and Greer 2022). Both usages of epigenetics maintain a core tenet that something distinct from the DNA sequence is inherited. Here I will use the definition of epigenetics in which specific modifications impacting gene expression, and subsequently phenotype, occur and are inherited.

Epigenetic modifications often change the way that DNA is packaged, altering its accessibility to factors that promote or inhibit transcription of RNA. Modifiers that are traditionally considered ‘epigenetic’ include DNA methylation (particularly cytosine methylation), RNA methylation, histone modifications (including methylation, acetylation, phosphorylation, ubiquitination, etc.), and nucleosome positioning, along with larger-scale changes to genome architecture (Handy et al. 2011; Mishra and Hawkins 2017). In addition, certain RNAs are themselves epigenetic, as they can affect their own transcription or transcription of other genes, and they can also impact post-transcriptional regulation of mRNA (*e.g*., small RNAs in RNA interference mechanisms) (Wei et al. 2017; Seroussi et al. 2022). Collectively, these epigenetic modifications each contribute to alterations in gene expression at the transcriptional or post-transcriptional level. However, notably, not all processes altering transcription or post-transcriptional regulation are epigenetic modifications (*e.g*., transcription factor binding).

Epigenetic modifications are inherited, but this can be either mitotically (across cell divisions in the soma) or meiotically (across generations via the germline). Mitotically-heritable epigenetic modifications are one mechanism by which unique cell types within the same individual are maintained, despite containing the same underlying DNA sequence (Almouzni and Cedar 2016). In addition, mitotically-heritable epigenetic states are common in cancer cells and play a role in cancer progression (Jones and Baylin 2002; Feinberg 2004; Baxter et al. 2014). Further, epigenetic modifications play a major role in phenotypic plasticity in response to particular environmental conditions (Kelly et al. 2012), and epigenetic mechanisms can maintain a plastic response in adulthood well after the initial trigger (Werner et al. 2023). In general, however, the epigenetic state of an organism’s cells, even if it is heritable mitotically and maintained throughout the lifespan, is not faithfully inherited meiotically through the germline, though there are some differences depending on the species. At one extreme, plant species including *Arabidopsis* have been observed to inherit cytosine methylation faithfully across meiosis for many generations, and differential methylation can be used to map quantitative traits to an ‘epigenetic quantitative trait locus’, which differs only in epigenetic modifications but not in underlying DNA sequence (Johannes et al. 2009; Cortijo et al. 2014; Kooke et al. 2015; Furci et al. 2019a; Liegard et al. 2019). Similarly, yeast have been shown to exhibit meiotically stable epigenetic states over many generations (Grewal and Klar 1996). For mammals, however, the notion of inheriting epigenetic modifications across generations has been more contentious, due to relatively few *bona fide* examples and the difficulty of studying mammals compared to other lab systems (Daxinger and Whitelaw 2012; Horsthemke 2018). In contrast, there are multiple examples of epigenetic inheritance in animal model systems including *Drosophila melanogaster* and *Caenorhabditis elegans* (Ruden and Lu 2008; Kelly 2014; Camacho et al. 2018; Moore et al. 2019; Baugh and Day 2020; Casas and Vavouri 2020; Evans et al. 2023). Notably, because *D. melanogaster* and *C. elegans* lack DNA cytosine methylation, mechanisms typically involve histone modifications and small RNAs.

The persistence of epigenetic modifications across generations that influence phenotype raises the possibility that epigenetic modifications can act like genetic variation, at least to some extent, in shaping the evolution of a population over time. Arguably, it is most interesting to investigate the role of epigenetic modifications in evolutionary trajectories when they act to some extent independently of genetic variation, acting as ‘drivers’ of phenotypic change rather than ‘effectors’ solely downstream of genetic variation (Sarkies 2020). That is, how do epigenetic modifications impact evolution when they arise spontaneously *a la* genetic mutations, and, while physically linked to genetic variants, impact fitness independently? This is distinct from asking how epigenetic modifications that arise because of environmental perturbations via plasticity (either within-generation or multigenerational plasticity) impact evolution. This is because plasticity can function like any other trait for which the genotype-phenotype map is of interest. Even in cases of multigenerational plasticity, it has been shown theoretically that heritable epigenetic responses to the environment can evolve given that the ancestral environment is predictive of the descendant environment (Herman et al. 2014). Epigenetic variants thar arise due to the environment (acting as ‘effectors’) can also impact evolutionary trajectories (subsequently acting as ‘drivers’), which has also been theoretically analyzed in several models (Geoghegan and Spencer 2012; Geoghegan and Spencer 2013).

Epimutation rates, used here to mean spontaneous epigenetic modifications (Le Goff et al. 2021), in contrast to plasticity-driven epigenetic marks, have been measured in a variety of species, and the consensus has consistently been that epimutations occur at higher rates than genetic mutations (Horsthemke 2006; Becker et al. 2011; Schmitz et al. 2011; Gravina et al. 2015; van der Graaf et al. 2015; Carja et al. 2017). Studies that have estimated epimutation rates tend to focus on CpG methylation, rather than other forms of epigenetic modifications. However, unlike germline genetic mutations that are faithfully inherited across generations, studies evaluating the inheritance patterns of epimutations without selection have been more limited, particularly in animals. Recently, experimental evolution was performed in *C. elegans* to analyze epimutation rates and the persistence of epimutations (Beltran et al. 2020). Results suggest that epimutations occur at higher rates than genetic mutations, can be maintained a few (2-10) generations without selection and exhibit a bias in the type of genes they target. Because it is known that genetic mutations exhibit a bias in terms of where they occur in the genome (which is often influenced by epigenetic factors) (Chen et al. 2012; Schuster-Bockler and Lehner 2012; Li et al. 2013; Li and Luscombe 2020; Monroe et al. 2022), a key functional difference between genetic mutations and epimutations is their duration of inheritance. It has been suggested that due to their limited duration, epimutations can only be maintained in populations long term if they offer a selective advantage (Beltran et al. 2020). Theoretical studies that have examined the inheritance of spontaneous epialleles have been relatively limited. Non-genetic inheritance has been broadly analyzed by extending the Price equation (Day and Bonduriansky 2011), and these results highlight the potential for the interaction of genetic and epigenetic alleles to affect evolutionary outcomes.

Here, we extend classic population genetic models to incorporate epimutations that impact fitness. Our results show that epialleles that would persist only a few generations without selection or ongoing epimutation can be maintained in long-term in epimutation-back-epimutation equilibrium. We show how this equilibrium changes when one epiallele has a selective advantage. Across genotypes, a compensatory or advantageous epiallele facilitates the maintenance of a deleterious genetic allele at a higher frequency than expected under mutation-selection balance. Collectively, these results suggest that a relatively simple accounting of epialleles into population genetic models can help explain both genetic and epigenetic variation in a population.

## MODEL

We consider two genetic alleles (*A*_1_ and *a*_2_) at a locus, each with an alternative epiallelic states (*A*_3_ corresponding to *A*_1_ and *a*_4_ corresponding to *a*_2_). Therefore, there are a total of four alleles at the locus, but only two underlying genotypes: *A*_1_ and *A*_3_ are the same genotype but can have distinct fitness values due to their epigenetic modifications, and a_2_ and a_4_ are also the same genotype but can have distinct fitness values. Each generation, the mutation rate, *μ*, is the rate at which *A*_1_ or *A*_3_ mutate to *a*_2_ or *a*_4_, respectively, and we assume there is no back-mutation. The epimutation rate, *ε*, is the rate of epimutation of *A*_1_ to *A*_3_ and *a*_2_ to *a*_4_. In this case, consistent with the limited inheritance potential of epimutations, the back-epimutation rate (*β*) models the rate at which *A*_3_ and *a*_4_ revert to *A*_1_ and *a*_2_. We consider diploid individuals in an infinite population with discrete generations. Mutation, epimutation, and back-epimutation occur in the gametes, while viability selection occurs in diploids. A schematic of the life cycle is shown in Figure 1A. Using *p*_1_(*t*), *p*_2_(*t*), *p*_3_(*t*), and *p*_4_(*t*) as the frequency of *A*_1_, *a*_2,_ *A*_3_, and *a*_4_ at time *t* in generations and representing the genotypic fitness values of the various diploid allelic combinations as *w*_11_, *w*_22_, *w*_33_, *w*_44_, *w*_12_, *w*_13_, *w*_14_, *w*_23_, *w*_24_, and *w*_34_, the recursion equations for each generation are given by:

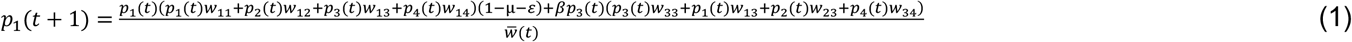

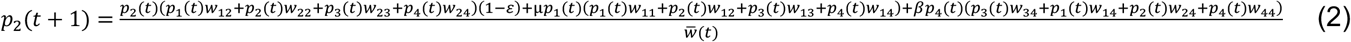

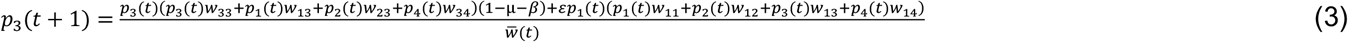

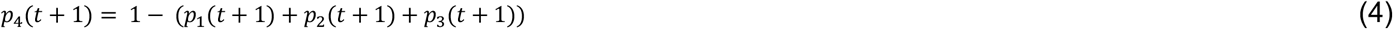

where 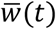 is the average fitness at time *t*. Transition probabilities from one epiallele to another, accounting for mutation, epimutation, and back-epimutation, are shown in Figure 1B. Under neutral conditions all fitness values are equal to 1, but when selection is occurring, each *w*_*nk*_ (for *n, k* equal to 1, 2, 3, or 4) can be written as 1-*s*_l_ for selection coefficient *s*_l_. For each fitness value, the corresponding selection coefficient can be written as *w*_11_ = 1-*s*_0_, *w*_22_ = 1-*s*_1_, *w*_33_ = 1-*s*_2_, *w*_44_ = 1-*s*_3_, *w*_12_ = 1-*s*_4_, *w*_13_ = 1-*s*_5_, *w*_14_ = 1-*s*_6_, *w*_23_ = 1-*s*_7_, *w*_24_ = 1-*s*_8_, and *w*_34_ = 1-*s*_9_.

**Figure 1:**
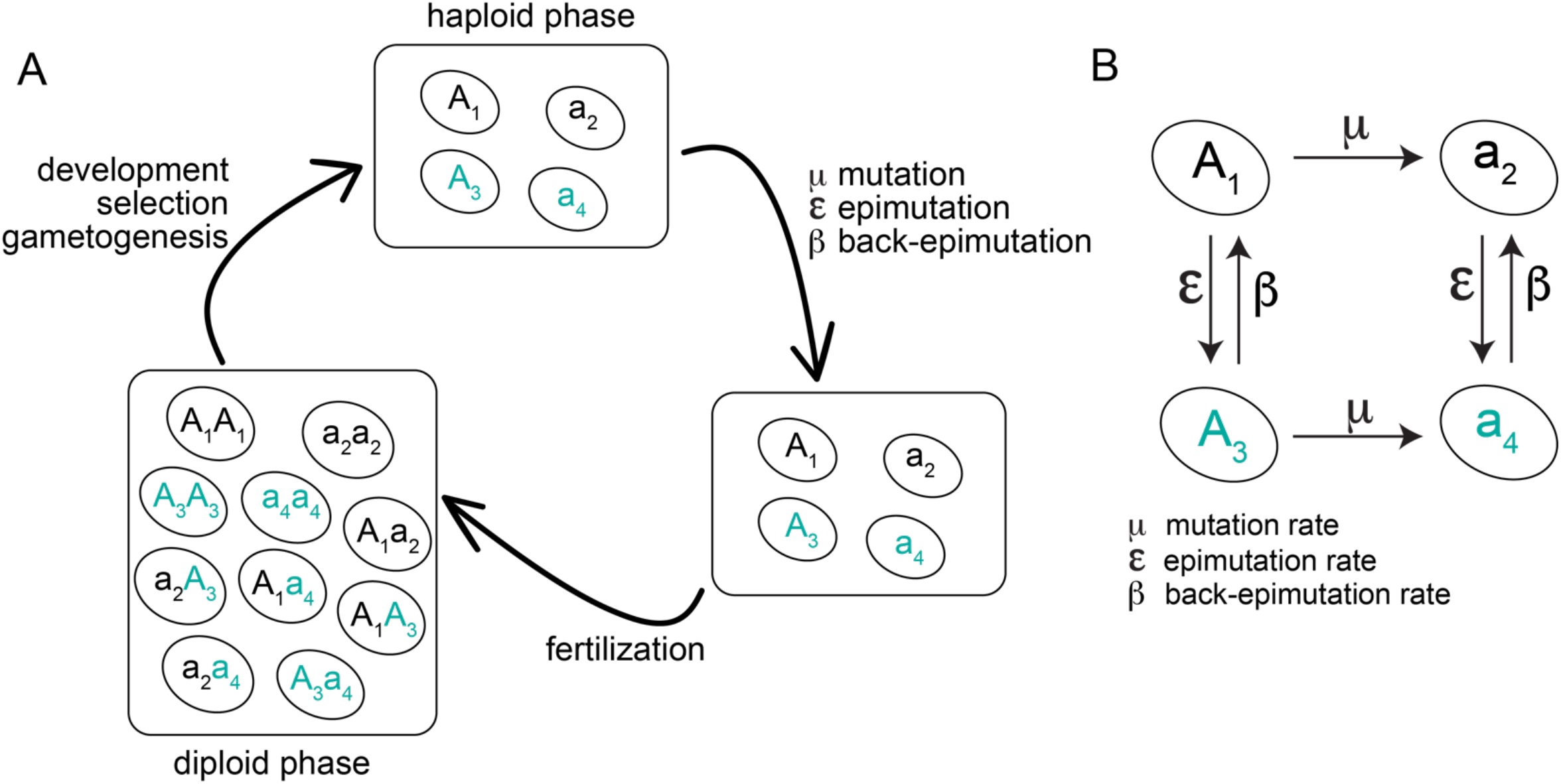
Life cycle diagram and transition probabilities for population epigenetic model. A. Four epialleles are possible: *A*_1_ and *a*_2_ are genetic alleles, while *A*_3_ and *a*_4_ represent alternative epiallelic states. Mutation, epimutation, and back-epimutation occur in the haploid gametes, and diploid combinations of each pair of epialleles are possible upon fertilization. Selection occurs on diploids, which then produce haploid gametes, completing the life cycle. B. Transition probabilities for mutation, epimutation, and back-epimutation for each type of epiallele to the other. Mutation occurs from *A*_1_ to *a*_2_ or from *A*_3_ to *a*_4_. Epimutation occurs from *A*_1_ to *A*_3_ or from *a*_2_ to *a*_4_. Finally, back-epimutation occurs from *A*_3_ to *A*_1_ or from *a*_4_ to *a*_2_. There is no parameter for back-mutation. Within a single generation, only one type of transition occurs, *i.e*., mutation, back-epimutation, and epimutation cannot happen sequentially or simultaneously in the same generation.

While epialleles are biologically distinct from genetic alleles, they are analogous to a multiallelic genetic model in which certain alleles have a high probability of ‘mutating’ (in our case, epimutating) to other, specific alleles. Equilibria for systems with *k* alleles were considered early in population genetics research (Kimura 1956; Mandel 1959), then subsequently expanded to include selection and/or mutation (Tallis 1966; Karlin 1980; Hadeler 1981; Varga and Zubiri 1993). These approaches have been based upon the assumption that mutation rates were low or equal across alleles, as is appropriate for approximately uniform changes in DNA sequence. The major difference with our analysis relative to other multiallelic models is that we allow epimutation and back-epimutations rates to be much higher than DNA-sequence mutation rates, which generates asymmetries in the equilibria that would not appear in sequence-based mutation approaches alone. We do make the simplifying assumption that multiple types of mutation do not occur in the same generation (*e.g*., it is not possible to have both mutation and back-epimutation occur at once).

We derive analytical solutions when possible, but a complete analysis requires numerical analysis in many cases. For numerical analysis, we use a mutation rate of 10^−7^ and epimutation rates ranging from 10^−5^ to 0.5. The rates used represent whole allele rates rather nucleotide-specific rates. We note that very low back-epimutation rates and very high epimutation rates are less likely to occur in nature. While epimutation rates are typically higher than mutation rates genome wide, the upper value we chose is likely significantly higher than a genome-wide rate. However, because there can be bias for which genomic loci experience epimutation, it is possible for the rate of epimutation to be higher than the genome-wide rate for a given allele (Beltran et al. 2020), and this high rate is meant to account for this circumstance. Finally, high back-epimutation rates are largely the parameter that distinguish a model of epigenetic inheritance from other models. While genetic mutations are unlikely to be truly reversible (and we assume that that is very unlikely here), epimutations are typically reversible on short timescales due to the rapid decay of epigenetic marks or modifiers. Based on the limited studies on the persistence of epimutations in animals, we have chosen two high back-epimutation rates of 0.33 and 0.1, which correspond to the maintenance of epialleles without selection for ∼3 or ∼10 generations, respectively (Sarkies 2020) (Figure 2). Based on the more extended persistence of epimutations observed in plants and yeast (Grewal and Klar 1996; Jacobsen and Meyerowitz 1997; Kakutani 2002), we have also included a back-epimutation rate of 0.01 (∼100 generations, Figure 2). Thus, we consider 0.33, 0.1, and 0.01 our ‘focal’ back-epimutation rates.

**Figure 2:**
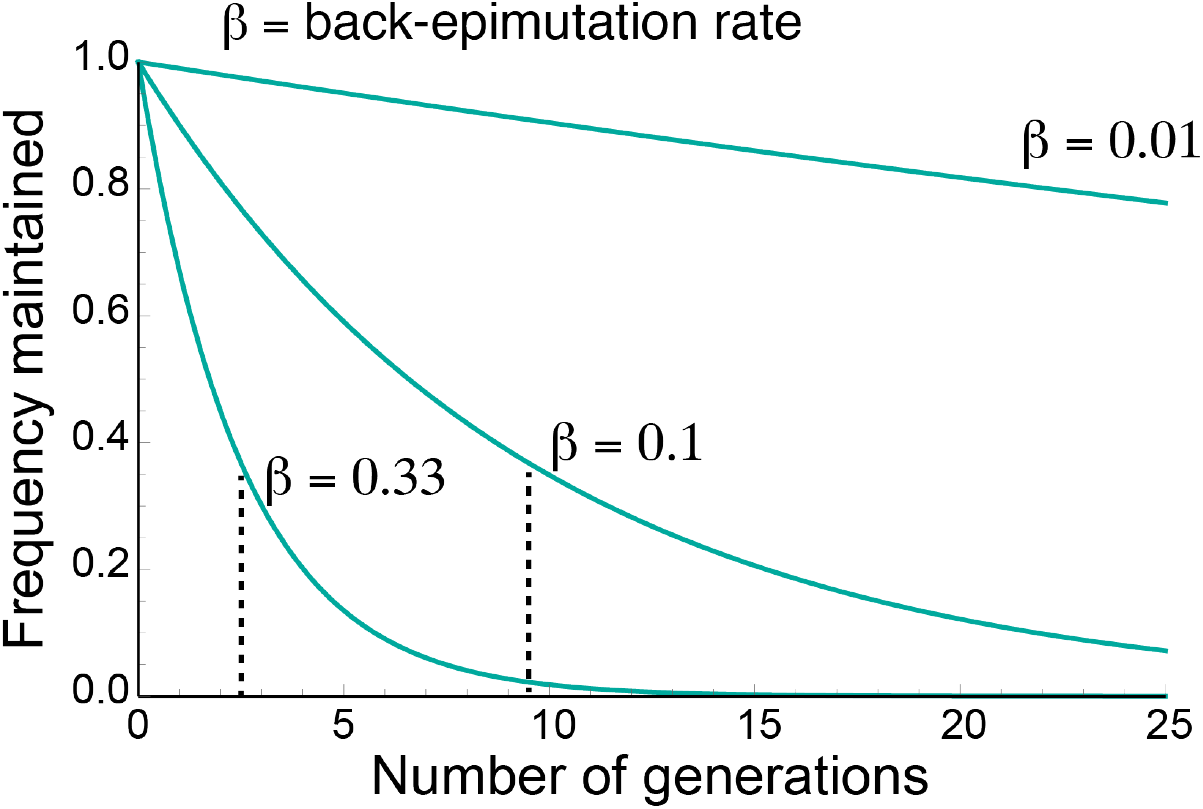
The back-epimutation rate (β) models the maintenance of an epiallele without selection. The back-epimutation rate models the maintenance of an epiallele without selection. In the absence of other factors, including new mutation, epimutation, or selection, the back-epimutation rate models the expected number of generations an alternative epiallele (*A*_3_ or *a*_4_) would be expected to be maintained. The equations plotted here are *p* = (1 – *β*)^t^, where *t* represents the number of generations, *β* is the back-epimutation rate, and *p* is the frequency of the epiallele that is maintained, which starts at 1. Examples are shown for three values of *β*: 0.33, 0.1, and 0.01, which model three different expected numbers of generations an epiallele would be maintained without selection: 2.5, 9.5, and 99.5, respectively. The expected value is indicated by the dotted line for *β* = 0.33 and 0.1.

## ANALYSIS METHODS

We used *Mathematica* version 12.2.0.0 to perform analysis of models and used R version 3.5.1 and Adobe Illustrator 2021 for data visualization. The Solve function was used to find solutions when a system could be solved analytically. For numerical solutions, equilibria were determined in *Mathematica* using the NSolve function by setting *p*_*x*_(t+1) = *p*_*x*_(t) in equations 1-3 above for a given parameter set, with *p*_4_ = 1 − (*p*_1_ + *p*_2_ + *p*_3_). For a given solution, working precision was set to 10 and solutions were restricted to be bounded by 0 and 1, inclusive.

Stability of equilibrium numerical solutions was determined by calculating the Jacobian matrix of the system via approximating the recursion equations above by the following continuous differential equations:

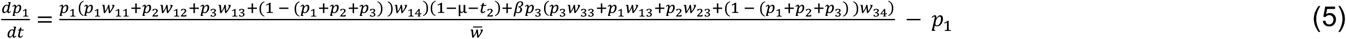

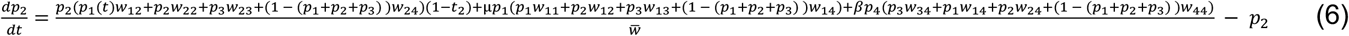

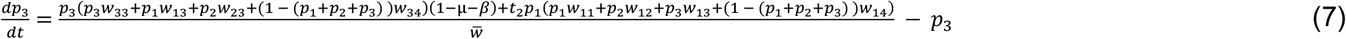

Allowing 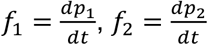, and 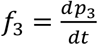, the Jacobian matrix is given by:

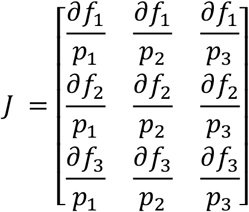

If all eigenvalues of this matrix were negative for a given parameter set, the equilibrium was considered stable; if at least one was positive, the equilibrium was considered unstable.

For all numerical analyses, the mutation rate was held constant at 10^−7^. We typically used fitness values of 1, 0.99, or 0.98 for values of each *w*_nk_ (for n,k equal to 1, 2, 3, or 4), but *w*_nk_ could lie anywhere between 0 and 1, inclusive, depending on the specific analysis and as indicated in each figure and in the text. For analyses that focused on the focal back-epimutation rates of 0.01, 0.1, and 0.33, epimutation rates were randomly drawn from a log-uniform distribution for rates between 10^−5^ and 0.5, then analyzed for each of the three back-epimutation rates. Results were exported into a .txt file, organized in Microsoft Excel as a .csv file, then analyzed in R. For these figures, solid and smooth lines were then drawn in Adobe Illustrator along points. Original graphs without lines drawn are available in Supplemental Figures 1-2. To analyze the full parameter space of epimutation and back-epimutation rates (*i.e*., the shaded red graphs in Figures 5-7), 300 random values were selected from a log-uniform distribution for both epimutation rates and back-epimutation rates between 10^−5^ and 0.5, then each combination of epimutation and back-epimutation rate (90,000 in total) was used to determine equilibrium solutions and stability. In rare cases, Mathematica failed to find equilibria for certain combinations of epimutation and back-epimutation rates. In other rare cases, the same equilibrium would be identified as an equilibrium multiple times for a given set of parameters. When these duplicates were identified, they were removed from the resulting data prior to plotting. If the source of an apparent irregularity could not be decisively determined, it was included in the final plot, such as some components of Figure 6F, Supplementary Figure 1B, and Supplementary Figure 2B.

## RESULTS

### Simplified models recapitulate standard population genetic solutions in the epigenetic context

Under neutral conditions (*w*_nk_ = 1 for all n,k), we find that both epialleles within a genotype (in this case, *a*_2_ and *a*_4_,) are maintained in long-term equilibrium in a manner that depends only on the epimutation and back-epimutation rates (Figure 3). Notably, this is exactly equivalent to genetic models that incorporate both forward and back genetic mutation (Wright 1931). Because back-mutation of a specific allele is rare, however, back-mutation is typically excluded from population genetic models that make simplifying assumptions. This results in the following relationship for the long-term equilibrium of *p*_4_, which we will refer to epimutation-back-epimutation equilibrium (EBE):

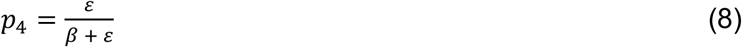

**Figure 3:**
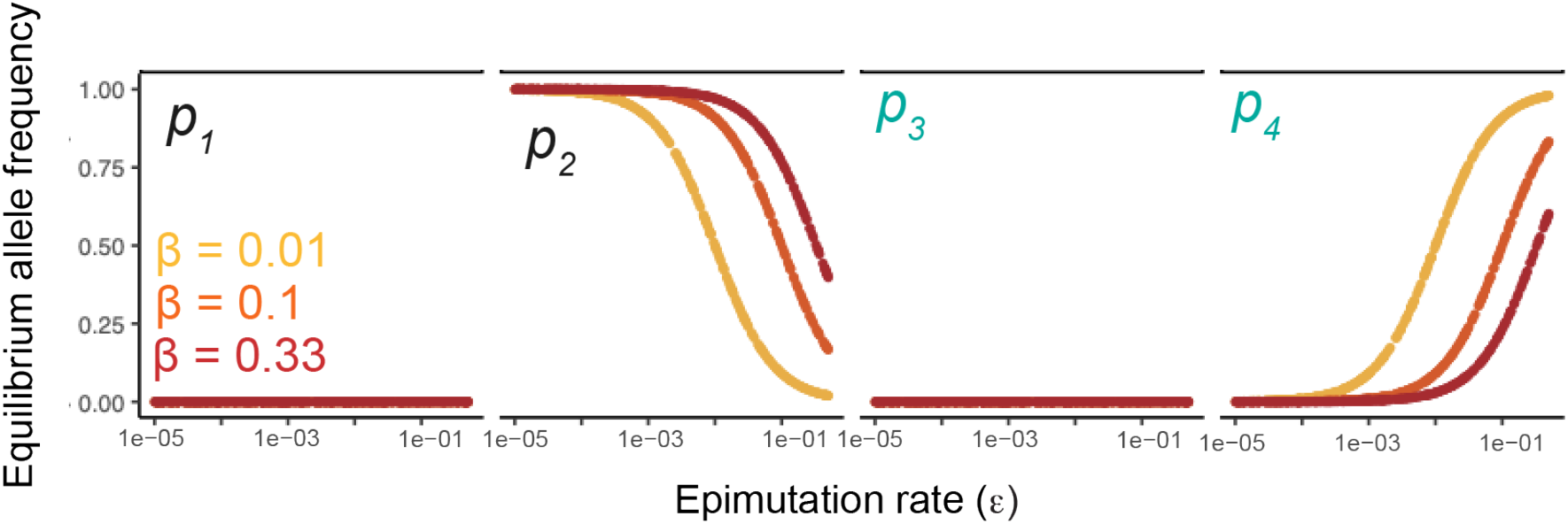
Epigenetic variation is maintained in epimutation-back-epimutation equilibrium under neutral conditions. When all fitness levels are set to 1, *a*_2_ and *a*_4_ alleles are found in equilibrium at frequencies that depend on the epimutation and back-epimutation rate, according to Equation 8.

Although biologically distinct from genetic mutations, epimutations should behave like genetic mutations as the back-epimutation rate (*β*) approaches zero. That is, we should be able to derive ‘epimutation-selection balance’ that is equivalent to mutation-selection balance in standard population genetic models. We focus again on *a*_2_ and *a*_4_, which are epialleles within the same genotype, and exclude the other genotype (*A*_1_ and *A*_3_). We thus have only three fitness values: *s*_1_, *s*_3_, and *s*_8_. We first solve equations for when *a*_2_ is dominant. Solving the recursion equations for the case when *β* = 0, *s*_1_ =0, and *s*_8_ = 0, yields the equilibrium frequency of *p*_4_:

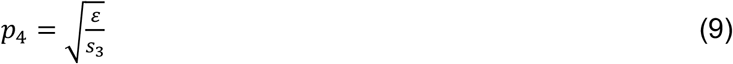

Similarly, incorporating a dominance coefficient, *h*, when *β* = 0, *s*_1_ =0, and *s*_8_ = 1-*hs*_3_ yields the equilibrium:

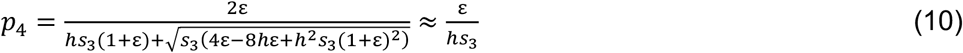

Equations 9 and 10 are likely biologically relevant for some plants, in which epialleles can be maintained long-term much like genetic mutations. Note that we can similarly obtain back-epimutation-selection balance if *ε* = 0, *s*_3_ =0, and *s*_8_ = 0, yielding the following equilibrium frequency of *p*_4_:

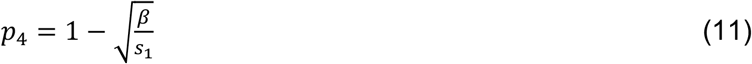

Because we typically expect epigenetic variants to be easily reversible, making *β* larger than the selection coefficient, in keeping with our intuition and in stark contrast to standard genetic mutation-selection balance, it is difficult to maintain an epigenetic allele unless it is also re-generated via epimutation.

### Compensatory epialleles cause deviations from mutation-selection balance by a common factor

Our main goal is to understand how all forces—epimutation, back-epimutation, mutation, and selection—interact in a system with four epialleles across two genotypes. We cannot analytically solve the full system with all selection coefficients as parameters, but we can solve cases that are direct extensions of classical mutation-selection balance. First, consider a case in which a reference genetic allele (*A*_1_) mutates to a deleterious recessive allele (*a*_2_) while the associated epialleles for each genotype, *A*_3_ and *a*_4_, are neutral with respect to *A*_1_. In other words, the *a*_4_ epiallele, while the same genotype as *a*_2_, has higher fitness than its counterpart. Biologically, this could model a common situation in which two genotypes each have an active and silent state (*e.g*. the locus is methylated or unmethylated). One genotype is deleterious when homozygous in its active state but is neutral with respect to the other alleles if at least one allele is silenced. The fitness of the other genotype does not depend on whether it is active or silenced. In our model this means all selection coefficients are 0 except for *s*_1_. Solving for the equilibrium for all four epialleles across both genotypes yields:

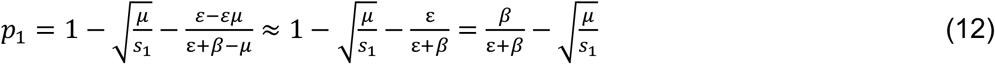

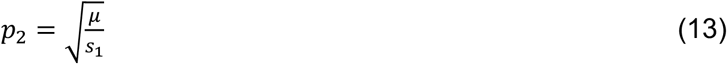

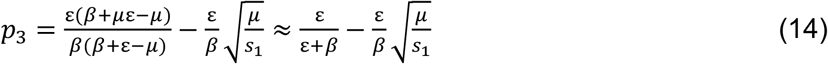

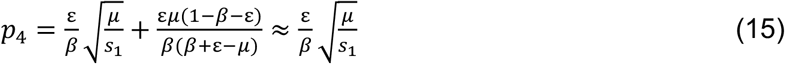

Recall that from a DNA sequence point of view, the observed allelic frequency within the population is the sum the of the modified and unmodified allelic states:

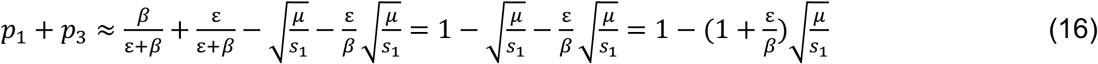

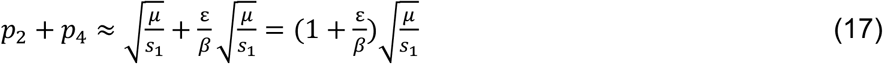

Under standard mutation-selection balance, we expect the dominant allele to be present at a frequency of 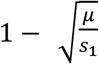, and the recessive allele to be present at frequency 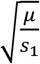. (This is also the case in the present model when *s*_1_, *s*_3_, and *s*_8_ all equal *s*_1_). However, in this case there is an alternative epigenetic state of the recessive deleterious allele that compensates for this deleterious effect. As a result, the equilibrium at a genotypic level is increased from mutation-selection balance by a factor of 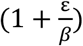. This quantity readily reveals the important factors affecting the maintenance of genetic variation. At one extreme, if *ε* > *β* we expect a large increase over mutation-selection balance, which could occur if new epigenetic variants are frequently generated and/or if they persist multiple generations. On the other hand, if new epigenetic variants arise infrequently and/or are readily reversed each generation (*ε* < *β*), then this will only minimally affect mutation-selection balance. And as expected, when no epigenetic variation is generated, the equilibria are exactly those derived under regular mutation-selection balance.

Alternatively, we can ask how these equilibria deviate from expectations from the EBE by summing modified and unmodified states across alleles:

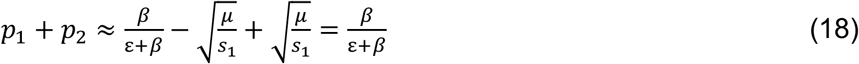

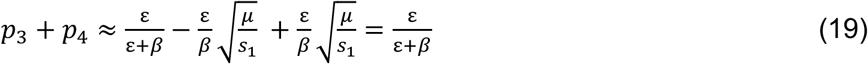

This shows there are no deviations from EBE by the introduction of a selection coefficient affecting only *w*_22_.

The analytical solution presented above applies in a case in which a deleterious mutation (*a*_2_) is completely recessive relative to the reference genetic allele (*A*_1_) as well as to alternative epialleles of either genotype (*A*_3_ or *a*_4_). However, it may be the case that *a*_2_ is not completely recessive, such as if silencing only one allele is not sufficient to mediate the negative consequences of mutation. Although we were unable to solve the system analytically when a general dominance coefficient (*h*) was incorporated, we can find a solution in the additive case when *h*=0.5. That is, *s*_4,_ *s*_7_, and *s*_8_ all equal *hs*_1_ when *h*=0.5 we find that solutions again deviate from mutation-selection balance – in this case 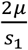 − by approximately a factor of 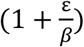. For the more general case of arbitrary dominance, we examined part of the parameter space numerically and examined whether our epigenetic frequency escalation term worked as a general approximation. We let *s*_1_ = 0.01 and *s*_4_, *s*_7_, *s*_8_ = *hs*_1_ and randomly selected values of *h* from 0 to 1. In all cases examined, the estimate of 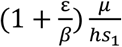 was a reasonable approximation for the sum of *p*_2_ and *p*_4_ and a better estimate than mutation-selection balance alone (Figure 4). Thus, these equations give us broad insight into how epigenetic variants that compensate from a deleterious mutation—whether that mutation is dominant, recessive, or with varying levels of dominance—cause higher equilibrium allele frequencies than those expected by mutation-selection balance alone, by deriving a fairly simple multiplier that is dependent on the relative magnitudes of *ε* and *β*.

**Figure 4:**
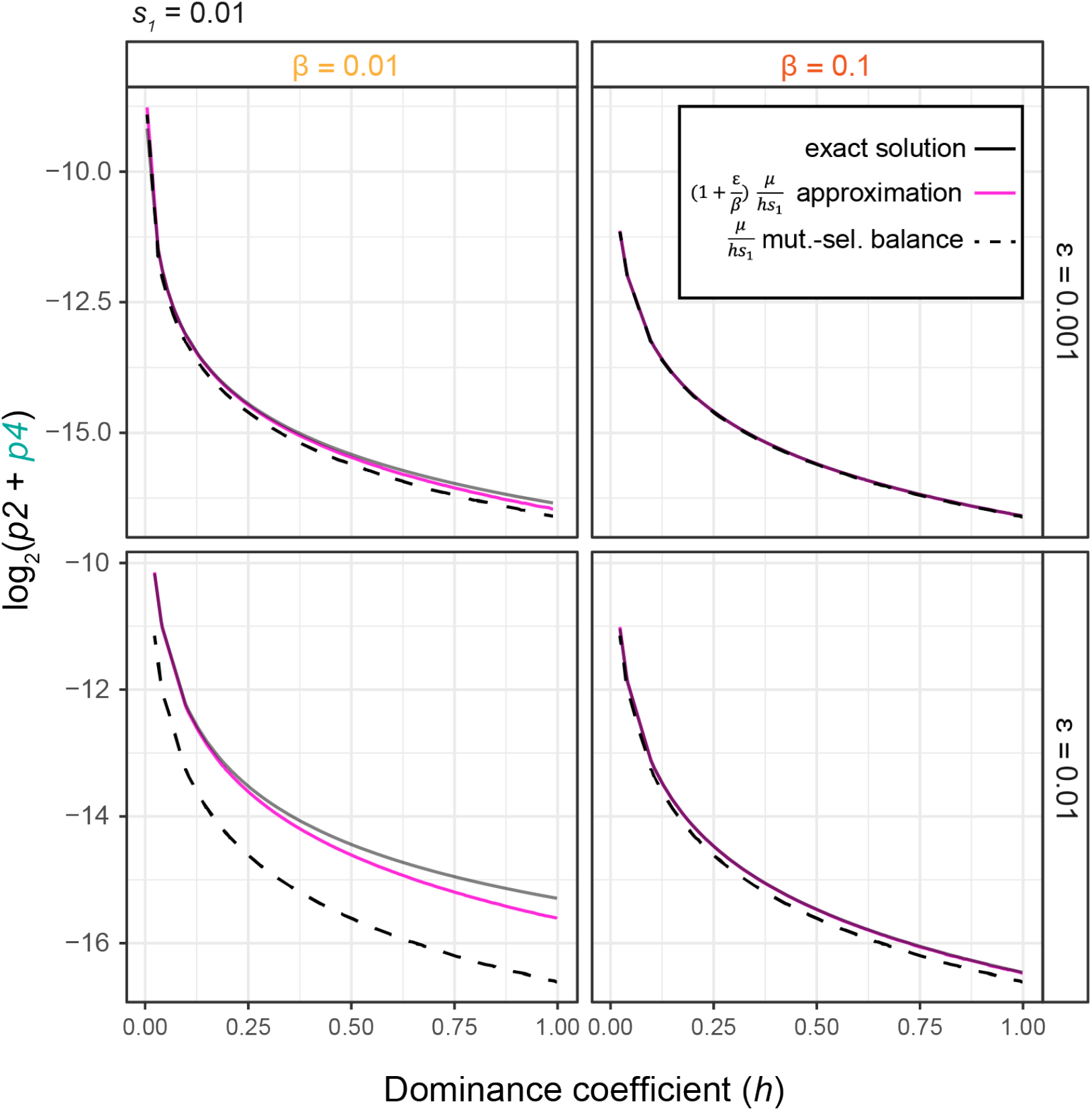
Numeric analysis suggests approximation with dominance coefficient accurately estimates epiallele equilibrium frequencies and deviation from mutation-selection balance Numerical solutions were obtained by solving the system of equations when *s*_*1*_=0.01, s_4_,s_7_,s_8_= *hs*_*1*_, *β* =0.01 or 0.1 and *ε*=0.001 or 0.01, for varying values of *h*. Exact numeric solutions (black) are shown alongside estimates based on the equation 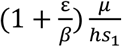 (pink) and estimates based on mutation-selection balance (dashed black).

### Epialleles under strong selection cause deviations from epimutation-back-epimutation equilibrium

We next asked how population dynamics are affected when a single epiallele yields a selective advantage relative to other alleles that are neutral with respect to each other, rather than compensating for a deleterious allele. While advantageous genetic alleles would be expected to be maintained at high frequency in a population, the high reversibility of epialleles makes it difficult to know how selection affects their long-term persistence. In our model we consider when the *a*_4_ epiallele has both a recessive and dominant fitness advantage relative to all other alleles. In the absence of any selective advantage, epialleles within a genotype will be in EBE, and by incorporating a selection coefficient of 0.01, our results show only mild deviations from EBE (Figure 5A-C). Throughout most of the parameter space, a single equilibrium exists, although in the recessive case there are sometimes two stable equilibria (Figure 5B). In the symmetric case when *A*_3_ has a selective advantage, we find that due to the asymmetry in the genetic mutation rate, it is possible for simultaneous maintenance of *A*_1_ and *a*_2_ to occur at low epimutation rates (Supplementary Figure 1), but for higher rates, patterns are consistent with results seen when *a*_4_ has a selective advantage. Notably, when *s*=0.2, *a*_4_ is maintained at high frequency even when the back-epimutation rate is high, highlighting the opportunity for an advantageous epiallele to significantly alter the dynamics of EBE. To systematically assess the role of selection coefficient on the dynamics of EBE, we randomly selected selection coefficients between 0 and 1 and allowed *a*_4_ to have a dominant or recessive selective advantage relative to all other alleles for specific epimutation and back-epimutation rates. In all cases, EBE is a good approximation for the equilibrium as long as the selection coefficient is small (less than 0.01), but it becomes an underestimate as the selection coefficient increases (greater than 0.1). EBE also is more likely to yield an accurate estimate if *ε* < *β*. Thus, population dynamics of small-effect variants that are inherited a limited number of generations – likely the dominant class of epigenetic variants in nature—are dominated by epimutation and back-epimutation rates and are well estimated by EBE. Nonetheless, large-effect epigenetic variants that are less readily reversible, if they arise, can significantly alter these dynamics.

**Figure 5:**
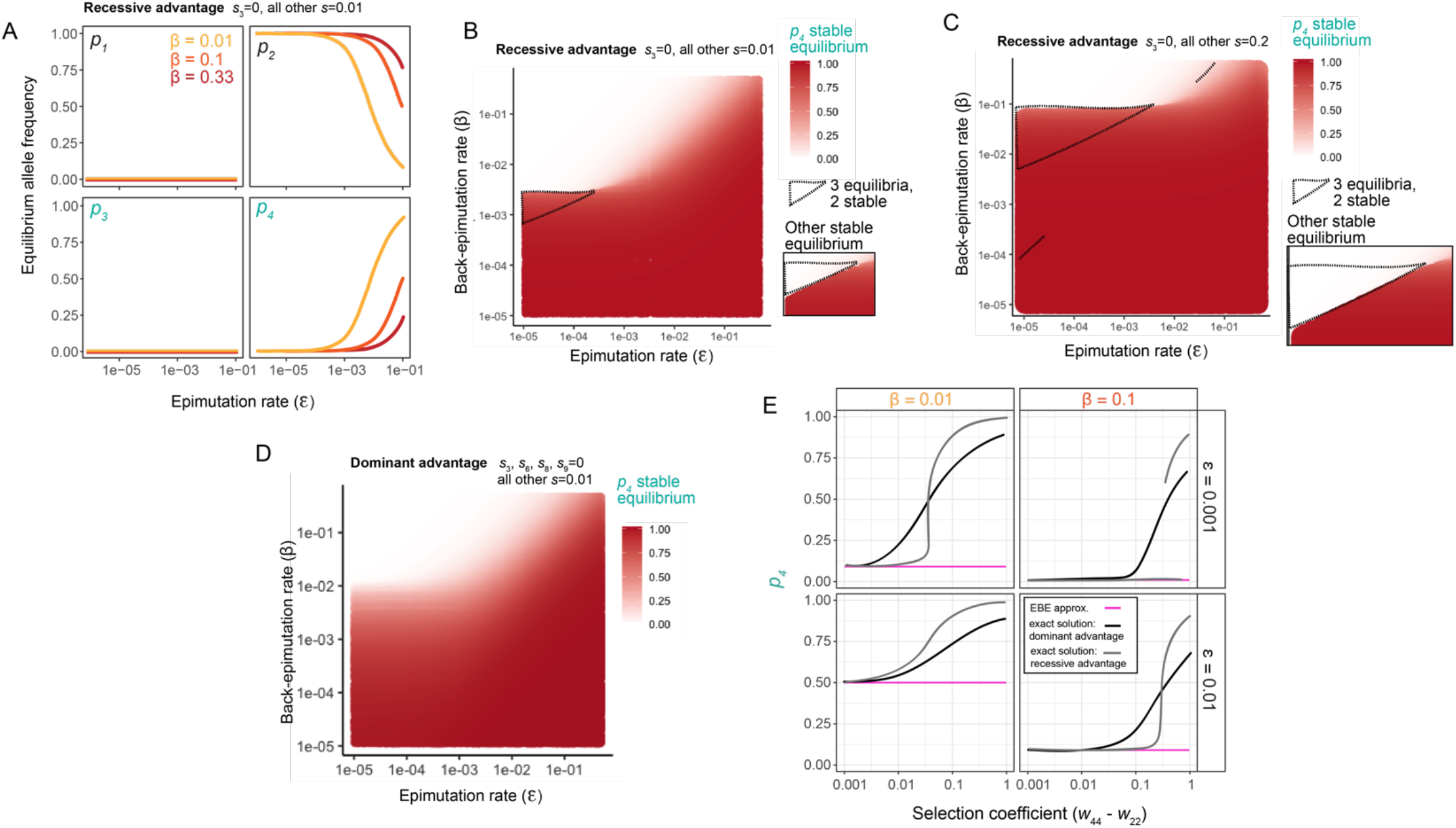
Strong selection drives deviations from epimutation-back-epimutation equilibrium A. Stable equilibrium allele frequency for *p*_*1*_ through *p*_*4*_ when *s*_3_=0 and other *s*=0.01 for back-epimutation rates of 0.01, 0.1, and 0.33. B-D. Stable equilibria for *p*_*4*_. In all cases, *p*_*1*_ and *p*_*3*_ are not maintained, and *p*_*2*_ = *1 – p*_*4*._ B. Stable equilibrium of *p*_*4*_ when *s*_3_=0 and other *s*=0.01 for a range of epimutation and back-epimutation rates. 3 equilibria are present within the dotted triangle (2 stable), and the other stable equilibrium is in the inset box. C. Same as (B) but other *s=*0.2. D. Stable equilibrium of *p*_*4*_ when *s*_3_, *s*_6_, *s*_8_, *s*_9_=0 and other *s*=0.01 for a range of epimutation and back-epimutation rates. E. Stable equilibrium of *p*_*4*_ when either *s*_3_=0 (recessive case) or *s*_3_, *s*_6_, *s*_8_, *s*_9_=0 (dominant case) and all other *s* vary between 0.001 and 1 for *β* =0.01 or 0.1 and *ε*=0.001 or 0.01.

### Certain advantageous epialleles maintain deleterious genetic alleles at high frequency

Thus far, we have examined population dynamics when an epiallele compensates for a deleterious allele (solution in equations 12-15) and when an epiallele has a selective advantage over other alleles that are neutral with respect to each other (Figure 5). Here, we combine these two cases and ask: how are population dynamics affected if an epiallele has higher fitness than both a reference and deleterious allele? For instance, methylated alleles could have higher fitness than an unmethylated alleles, regardless of the underlying DNA sequence, while the fitness of an allele when it is unmethylated is dependent upon its genetic sequence. For comparison’s sake, we first show the case of an epiallele compensating for a recessive deleterious allele numerically when *s*_1_ = 0.01 (Figure 6A-B). We then show how these dynamics change when, instead of merely compensating for a deleterious allele, *A*_3_ or *a*_4_ epialleles have a selective advantage compared to other alleles in a dominant manner (Figures 6C-D). This leaves *A*_1_*A*_1_, *A*_1_*a*_2_, and *a*_2_*a*_2_ with intermediate fitness and *a*_2_*a*_2_ alone with the lowest fitness. The primary difference in the resulting equilibria between these two cases is for *A*_3_ and *a*_4_ equilibria when *ε* and *β* are both low. Thus, for high *β*, equations 12-15 are a good approximation of equilibria. However, these results change substantially if only *a*_4_ has a dominant selective advantage, but *A*_3_ does not (Figure 6E-F). Here, *s*_1_=0.02, *s*_3_, *s*_6_, *s*_8_, *s*_9_ = 0, and all other *s*=0.01. We find that the frequency of *a*_2_, despite being recessive and deleterious, increases to high frequency, well beyond the expectation of mutation-selection balance. All four epialleles can be maintained at appreciable frequency simultaneously, which is unique among situations considered thus far. In this case, the driving factor is likely that *a*_2_ and *a*_4_ – epialleles within a genotype – correspond to the extremes of the fitness distribution. This maintenance of all four epialleles across both genotypes also occurs if we look at the symmetric case in which diploids containing *A*_1_ and *A*_3_ are at the extremes of the fitness distribution (Supplementary Figure 2), highlighting that this phenomenon is not merely a function of the unidirectional mutation rate.

**Figure 6:**
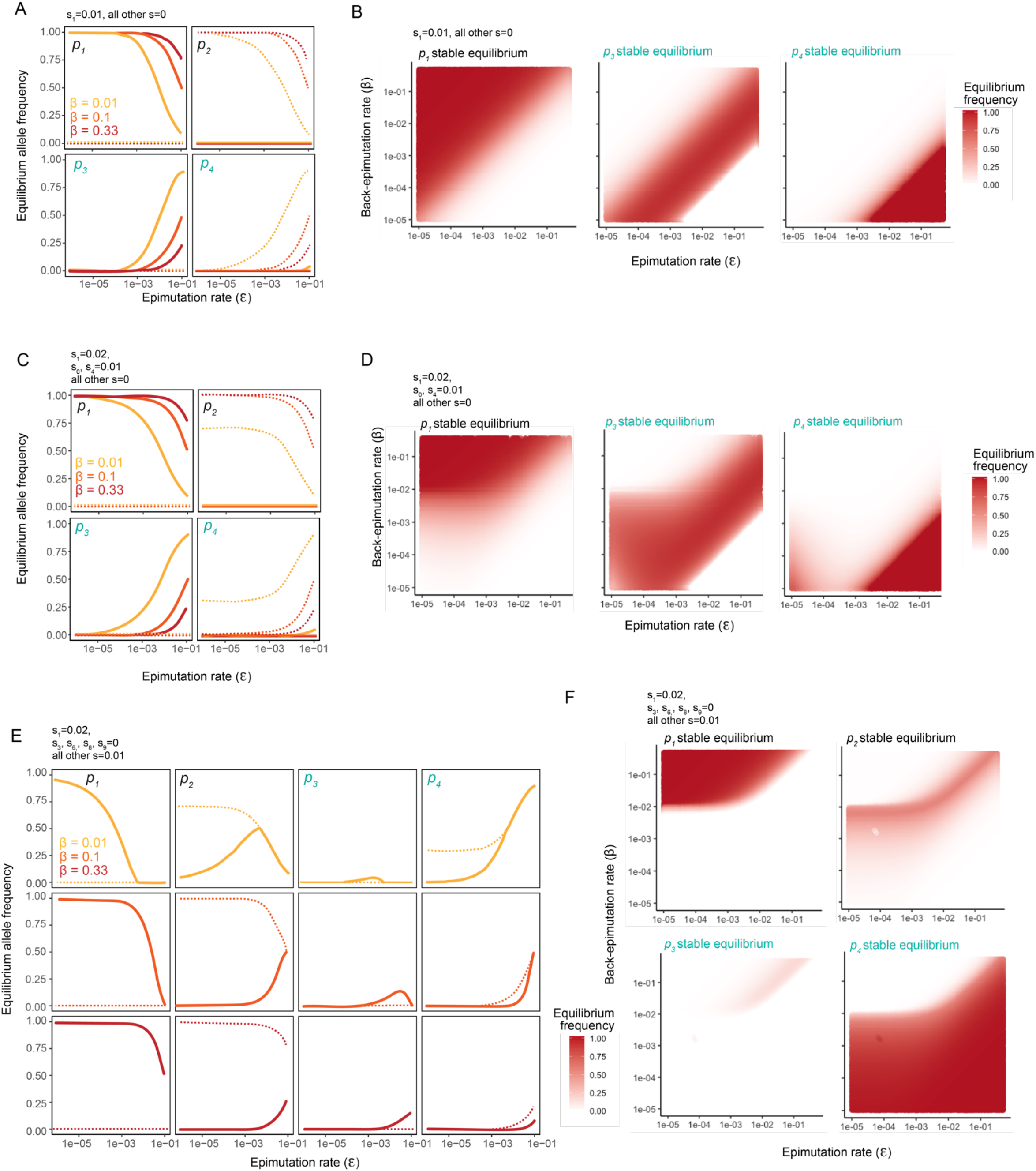
Epialleles compensating for deleterious recessive alleles drive deleterious alleles to high frequency A. Stable equilibrium allele frequencies (solid lines) and unstable equilibrium allele frequencies (dashed lines) when *s*_1_=0.01 and all other *s*=0, for back-epimutation rates of 0.01, 0.1, and 0.33. B. Stable equilibrium when *s*_1_=0.01 and all other *s*=0 for a range of epimutation and back-epimutation rates. *p*_*2*_ is not shown because it is very close to 0 and not visible at the scale shown. C. Stable equilibrium allele frequencies (solid lines) and unstable equilibrium allele frequencies (dashed lines) when *s*_1_=0.02, *s*_0_,*s*_4_=0.01, and all other *s*=0 for back-epimutation rates of 0.01, 0.1, and 0.33. D. Stable equilibrium when *s*_1_=0.02, *s*_0_,*s*_4_=0.01, and all other *s*=0 for a range of epimutation and back-epimutation rates. *p*_*2*_ is not shown because it is close to 0. E. Stable equilibrium allele frequencies (solid lines) and unstable equilibrium allele frequencies (dashed lines) when *s*_1_=0.02, *s*_3_,*s*_6_,*s*_8_,*s*_9_=0, and all other *s*=0.01 for back-epimutation rates of 0.01, 0.1, and 0.33. F. Stable equilibrium when *s*_1_=0.02, *s*_3_,*s*_6_,*s*_8_,*s*_9_=0, and all other *s*=0.01 for a range of epimutation and back-epimutation rates.

When the deleterious allele is instead dominant, and the compensatory epiallele is recessive, we expect the genetic allele to be maintained at a frequency of 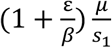. We show this result numerically in Figure 7A-B for *s*_1_, *s*_4_, *s*_7_, *s*_8_ =0.01 when all other *s*=0. When only diploids containing epialleles *A*_3_ and *a*_4_ have the highest fitness (*s*_2_,*s*_3_,*s*_9_=0), and those containing *A*_1_ are intermediate, we find differences primarily in the equilibria of *A*_3_ and *a*_4_ when *ε* and *β* are both low (Figure 7C-D). This is analogous to Figure 6 for the recessive case and our derived deviation from mutation-selection balance, 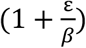, gives a reasonable approximation of equilibria for high *β*. An exception again occurs when a single epiallele exhibits a recessive advantage relative to all other alleles (Figure 7E-F). Here, we find multiple equilibria even for high *β*, and one of these equilibria includes the maintenance of the deleterious *a*_2_ at high frequency. Unlike the recessive case shown in Figure 6E-F, all four epialleles are not simultaneously present at high frequency. Instead, the mix of epialleles consists primarily of *A*_1_ and *A*_3_ or *a*_2_ and *a*_4_. Nonetheless, it is striking that a recessive advantageous epiallele can potentially cause such deviations from expectation for a deleterious dominant mutation. As before, this is likely due to the epialleles within a genotype corresponding to the extremes of the fitness distribution.

**Figure 7:**
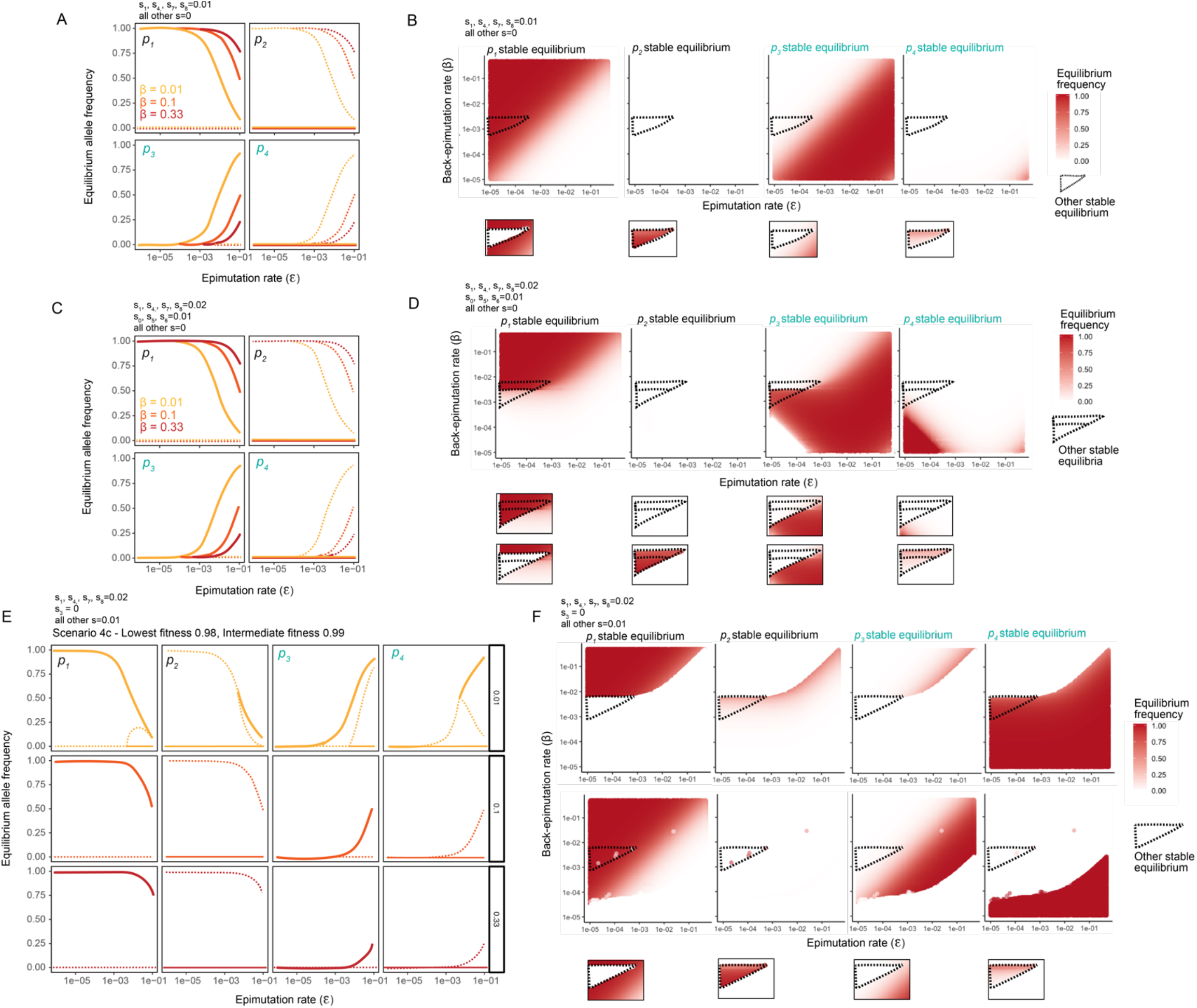
Epialleles compensating for deleterious dominant alleles drive deleterious alleles to high frequency A. Stable equilibrium allele frequencies (solid lines) and unstable equilibrium allele frequencies (dashed lines) when *s*_1_, *s*_4_, *s*_7_, *s*_8_ = 0.01 and all other *s* = 0 for back-epimutation rates of 0.01, 0.1, and 0.33. B. Stable equilibria when *s*_1_, *s*_4_, *s*_7_, *s*_8_ = 0.01 and all other *s* = 0 for a range of epimutation and back-epimutation rates. Within the dotted triangle, a second stable equilibrium is present. C. Stable equilibrium allele frequencies (solid lines) and unstable equilibrium allele frequencies (dashed lines) when *s*_1_, *s*_4_, *s*_7_, *s*_8_ = 0.02, *s*_0_, *s*_5_, *s*_6_ = 0.01, all other *s* = 0, and back-epimutation rates are 0.01, 0.1, and 0.33. D. Stable equilibria when *s*_1_, *s*_4_, *s*_7_, *s*_8_ = 0.02, *s*_0_, *s*_5_, *s*_6_ = 0.01, and all other *s* = 0 for a range of epimutation and back-epimutation rates. Within the smallest dotted triangle, 3 stable equilibria are present. Within the larger dotted triangle but excluding the smaller triangle, 2 stable equilibria are present. E. Stable equilibria allele frequencies (solid lines) and unstable equilibrium allele frequencies (dashed lines) when *s*_1_, *s*_4_, *s*_7_, *s*_8_ = 0.02, *s*_3_ = 0, all other *s* = 0.01, and back-epimutation rates are 0.01, 0.1, and 0.33. F. Stable equilibria when *s*_1_, *s*_4_, *s*_7_, *s*_8_ = 0.02, *s*_3_ = 0, and all other *s* = 0.01 for a range of epimutation and back-epimutation rates. The full parameter space is shown for two stable equilibria (each row is a different equilibrium) because both are present throughout much of the space. A third stable equilibrium is found within the dotted triangle.

Because we twice observed that maintaining a deleterious allele (*a*_2_) at high frequency is dependent on the fitness advantage of its corresponding epiallele (*a*_4_), we wanted to further assess the parameters for which this is possible. When *a*_2_ is recessive deleterious and *a*_4_ is dominant advantageous, *a*_2_ is maintained at high frequency alongside all other epialleles simultaneously (Figure 6E-F, Figure 8A). When *a*_2_ is dominant deleterious and *a*_4_ is recessive advantageous, we find that *a*_2_ can be maintained at high frequency but only alongside *a*_4_ (Figure 7E-F, Figure 8A). When the alternative epiallele is deleterious, this dynamic is reversed (Figure 8A). That is, when *a*_2_ is recessive advantageous and a_4_ is dominant deleterious, only *a*_2_ and *a*_4_ are maintained at appreciable frequency, but when *a*_2_ is dominant advantageous and *a*_4_ is recessive deleterious, all four epialleles are maintained (Figure 8A). In both cases for which all four epialleles are maintained, one epiallele acts dominantly to increase fitness, while its corresponding epiallele within the same genotype has the lowest fitness and acts in a recessive manner (and the symmetric situations for *A*_3_ and *A*_1_ also show these same dynamics; see Supplementary Figure 2). In contrast, when deleterious alleles are maintained, but not in a manner that promotes multiple genetic alleles simultaneously, this is due to a recessive fitness advantage and dominant deleterious fitness. Because deleterious genetic variation can be maintained in long-term equilibrium alongside the favored allele, we assessed numerically how the fitness of the deleterious allele affects these dynamics. We held the epimutation rate constant (at either 10^−5^ and 0.01) but allowed the fitness of *a*_2_*a*_2_ (*w*_*22*_) to vary. While the magnitude of the equilibrium frequency for *a*_2_ varies depending on the fitness, the parameters (particularly, the back-epimutation rate) for which *a*_2_ is maintained do not (Figure 8B-C). As a result, in order to maintain deleterious genetic variation when the back-epimutation rate is high (0.1 or greater), the epimutation rate must also be high, and this is true regardless of the fitness of the deleterious allele.

**Figure 8:**
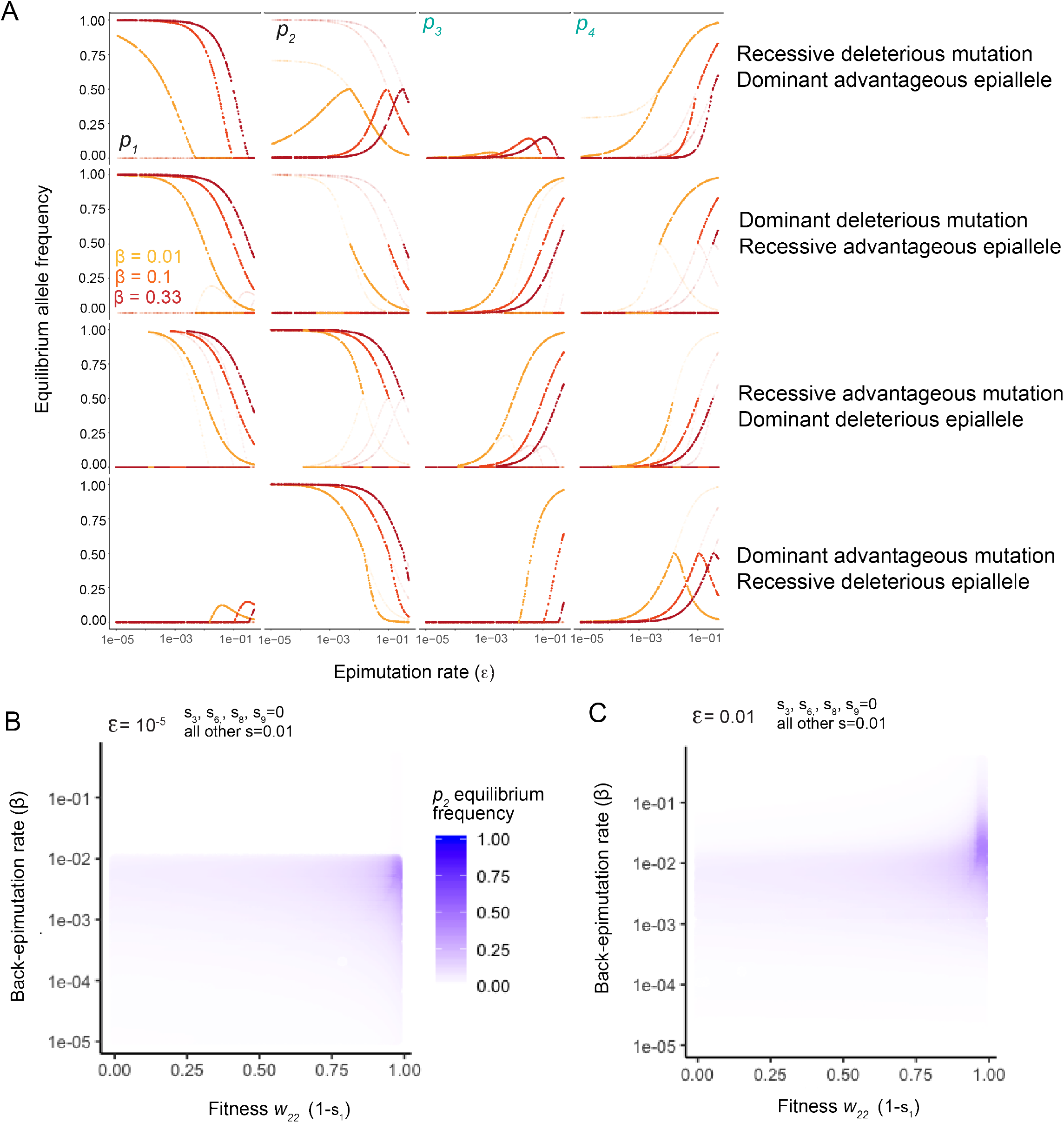
Deviations from mutation-selection balance depend on dominance status of mutations and compensatory epialleles A. Equilibrium frequencies for *p*_1_ through *p*_4_. Opaque points represent stable equilibria, while faded points are unstable equilibria. B-C. *w*_*22*_ (the lowest fitness) was allowed to take any value between 0 and 0.98 and back-epimutation rate was allowed to take any value between 10^−5^ and 0.5. The long-term equilibrium of *p*_*2*_ (the frequency of *a*_2_) is indicated by the shading of blue. B. The epimutation rate is held constant at 10^−5^. C. The epimutation rate is held constant at 0.01.

## DISCUSSION

Here we developed and analyzed an extension of a population genetic model that incorporates epigenetic alleles. We solved relevant simplifications of the full system analytically and special cases of the full system numerically. First, the neutral case (all *w*_nk_ = 1) shows that multiple epigenetic states within a single genotype can be maintained in a manner that depends only on epimutation and back-epimutation rates, called epimutation-back-epimutation equilibrium (EBE). The neutral case is directly analogous to the maintenance of genetic variation by mutation and back-mutation, although epimutation and back-epimutation occur at much higher rates than genetic mutation in nature and thus can drive substantial epigenetic variation. Second, we show that when the back-epimutation rate is 0 (meaning epigenetic variants are not reversible and thus act similarly to genetic mutations) and selection acts against new epiallales, we derive the equation for epimutation-selection balance that is directly analogous to mutation-selection balance. Third, we show that when an epimutation compensates for a recessive or dominant deleterious genetic allele, we find deviations from mutation-selection balance by a common factor, 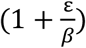. Finally, we use numerical analysis to analyze specific cases of the full model when both genetic and epigenetic variation are present. Of particular interest, we find instances of dramatic deviation from mutation-selection balance when the two epialleles of the same genotype are at the extremes of the fitness distribution. Collectively, this demonstrates the importance of considering epigenetic variation to understand evolutionary dynamics of populations.

The incorporation of epigenetic alleles in population genetic models has important implications for the maintenance of epigenetic and genetic variation. First, the maintenance of *epigenetic* variation occurs in all situations examined and is approximated well by epimutation-back-epimutation equilibrium when selection coefficients are small. This suggests the maintenance of epigenetic variation may be relatively common if the epimutation rate is sufficiently high. Second, the maintenance of *genetic* variation occurs when a deleterious allele is compensated for by an alternative epiallele. This likely does not represent a large deviation in most cases, but it highlights the possibility for large perturbations that may occur for high *ε* and low *β*. We discuss these major results below. In addition, we consider the potential role of two factors not considered here -- drift and the environment.

### The maintenance of epigenetic variation may be common

Our results suggest that when an epigenetic variant arises, it may be maintained in long-term stable equilibrium even if it is highly reversible (*i.e*., even if it has a high back-epimutation rate). This is true regardless of whether mutation and epimutation of an allele result in similar or distinct fitness effects. However, it is notable that for highly reversible epimutations (those maintained 3 to 10 generations without selection) to be maintained long-term at appreciable frequency in the absence of strong selection, they must also be generated at relatively high frequency (∼0.001 to 0.01 probability each generation). While epimutations occur at higher frequency than genetic mutations, this range is higher than expected under non-stressful, baseline conditions and when considering a genome-wide rate. As a result, it will be important for future empirical work to determine 1) whether there are environmental conditions for which the epimutation rate may be higher, and 2) whether there are epimutation hotspots in the genome for which epimutation rates are higher. It has been observed that certain stresses can act to increase phenotypic variation, possibly due to increasing epigenetic variation, and transgenerational epigenetic effects have also been identified in response to stress. In addition, the observation that transgenerational effects are more robust if individuals are exposed to environmental stressors for multiple generations lends credence to the idea the epimutation rates may be higher under stress (Vogt and Hobert 2023). However, to our knowledge, experimental evolution in an ongoing stressful environment with monitoring of epimutations across generations has not been performed. In addition, the potential role of drift remains unclear, though the maintenance of epialleles within a given population is likely to largely depend on the population size, which can be explored in future theoretical studies.

### Design of experiments to assess the maintenance of epigenetic variation

In measurements of epigenetic regulation across individuals in a population, it is often unclear whether differences across individuals are due to technical variation, ‘random’ noise, or meaningful biological differences. Our equation approximating the maintenance of an epiallele (Equation 8) may be a useful consideration when attempting to differentiate these possibilities. Epimutation rates can be empirically measured, as can their reversibility. Given these two parameters, it is straightforward to calculate how much variation is expected to be present at a given locus. On the other hand, these parameters allow for more precise experimental evolution paradigms to proceed. If epimutation rates are very low, then epigenetic variation is likely not to be maintained, assuming that the environment is constant and the population is sufficiently large for the present model to be applicable.

### Compensatory epimutations cause predictable deviations from mutation-selection balance

In standard population genetics, the equilibrium of a deleterious recessive or dominant allele is well known. Here, we introduce an epiallele of a deleterious allele that compensates for this effect and find that the genotype containing both the deleterious and compensatory epialleles deviates from the expectation of mutation-selection balance by a factor of 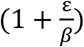. Thus, in cases in which epimutation and back-epimutation are present and afford some degree of compensation for a deleterious mutation, mutation-selection balance will be an underestimate of the deleterious allele in the population. It will likely be difficult to assess whether epimutation and back-epimutation are playing a role in natural populations, given that these parameters may depend on an environment that will be difficult to recapitulate in the lab. However, at least one clue that these may be playing a role is if significant variation in the phenotype is present for a deleterious allele. For example, if the average fitness of an alternative genotype is lower than the reference genotype, but there is significant variability across individuals in the alternative genotype within a single environment, this may indicate that cryptic epialleles exist within the genotype. Directly associating an epigenetic mark, such as cytosine or histone methylation, with a phenotype can provide support to this idea in this case.

### The maintenance of deleterious genetic variation at high frequency is possible but likely rare

While epialleles were maintained for certain parameters in every situation analyzed, deleterious genetic variation was typically not maintained at high frequency. Sometimes, we find that *A*_3_ and *a*_4_ are maintained together at high frequency (see the boundary between *A*_3_ and *a*_4_ in Figures 6 and 7), but this is in a part of the parameter space for which the back-epimutation rate is very low, causing the epimutation to behave more like a genetic mutation. In these cases, it is relatively unsurprising that an advantageous and minimally-reversible epiallele is maintained. Such a scenario might be applicable when a gene silencing state via DNA methylation is required for fitness, and the silenced state is inherited; in this case, if a genetic mutation occurs, the mutation may not realize its effect. However, in the context of highly reversible epigenetic variants, deleterious variation can only be maintained at high frequency when the deleterious allele has an alternative epiallele with higher fitness than other alleles in the system (see Figures 6E-F, Figure 7E-F, Figure 8). In both cases, the extremes of the fitness distribution include the alternative epialleles of *a* – with *a*_2_*a*_2_ having the lowest fitness and *a*_4_*a*_4_ having the highest fitness. While it is not difficult to imagine that an epiallele could compensate for a deleterious genetic allele resulting in the deviation from mutation-selection balance discussed above, it is more difficult to imagine that an epimutation would make the deleterious allele *more* fit than the non-deleterious allele. However, again, the role of the environment and drift were not considered in the present study, and analyzing both of these possibilities could lend additional insight into the feasibility of such a scenario in nature. For example, if a deleterious allele is only deleterious in a given environment, is it possible that occasionally encountering a more favorable environment can facilitate maintenance of the deleterious allele? In addition, is it possible that deleterious variation will be maintained if the population is sufficiently small? Such questions can be answered by extensions to the present model and will inform future empirical studies.

## CONCLUSIONS

Understanding the forces that maintain variation within natural populations represents a very long term and central part of evolutionary genetics (Lewontin 1974; Kimura 1983; Marion and Noor 2022). One of the earliest observations in the field is that there is far more variation at the molecular level than would be expected via purifying selection. Our results demonstrate that epigenetic modifications can be added to the long list of other potential factors such as balancing selection that can elevate allele frequencies above the mutation-selection balance expectation. There has long been interest in the potential for non-genetic inheritance to have an influence on evolutionary outcomes (Kirkpatrick and Lande 1989; Mousseau and Fox 1998; Jablonka and Raz 2009; Richards et al. 2012). The last decade has led to rapid increase in knowledge of the molecular underpinnings of epigenetic variation and the inheritance of these effects for a number of generations (Silveira et al. 2013; Öst et al. 2014; Schott et al. 2014; Siklenka et al. 2015; Ciabrelli et al. 2017; Klosin et al. 2017; Yu et al. 2018; Furci et al. 2019b; Moore et al. 2019; Kaletsky et al. 2020; Beck et al. 2021). While the pace of discovery on the molecular side is accelerating, the population genetic implications of these mechanisms have yet to be fully explored. Here, we examine a case in which epimutations are “random” with respect to any induction by the environment. In this case, variation is generated at the “phenotypic” level via the stable maintenance epialleles, and in some cases at the genetic level via the maintenance of deleterious alleles that whose effects would otherwise appear cryptic in the absence of specific information about potential epigenetic modification. Indeed, these epigenetic effects exist in the liminal space between genotype and phenotype in the classical models of genetic inheritance, and, depending on their prevalence, hold the potential to further complicate the already complex landscape functional variation within natural populations.

## SUPPLEMENTARY FIGURES

**Supplementary Figure 1:**
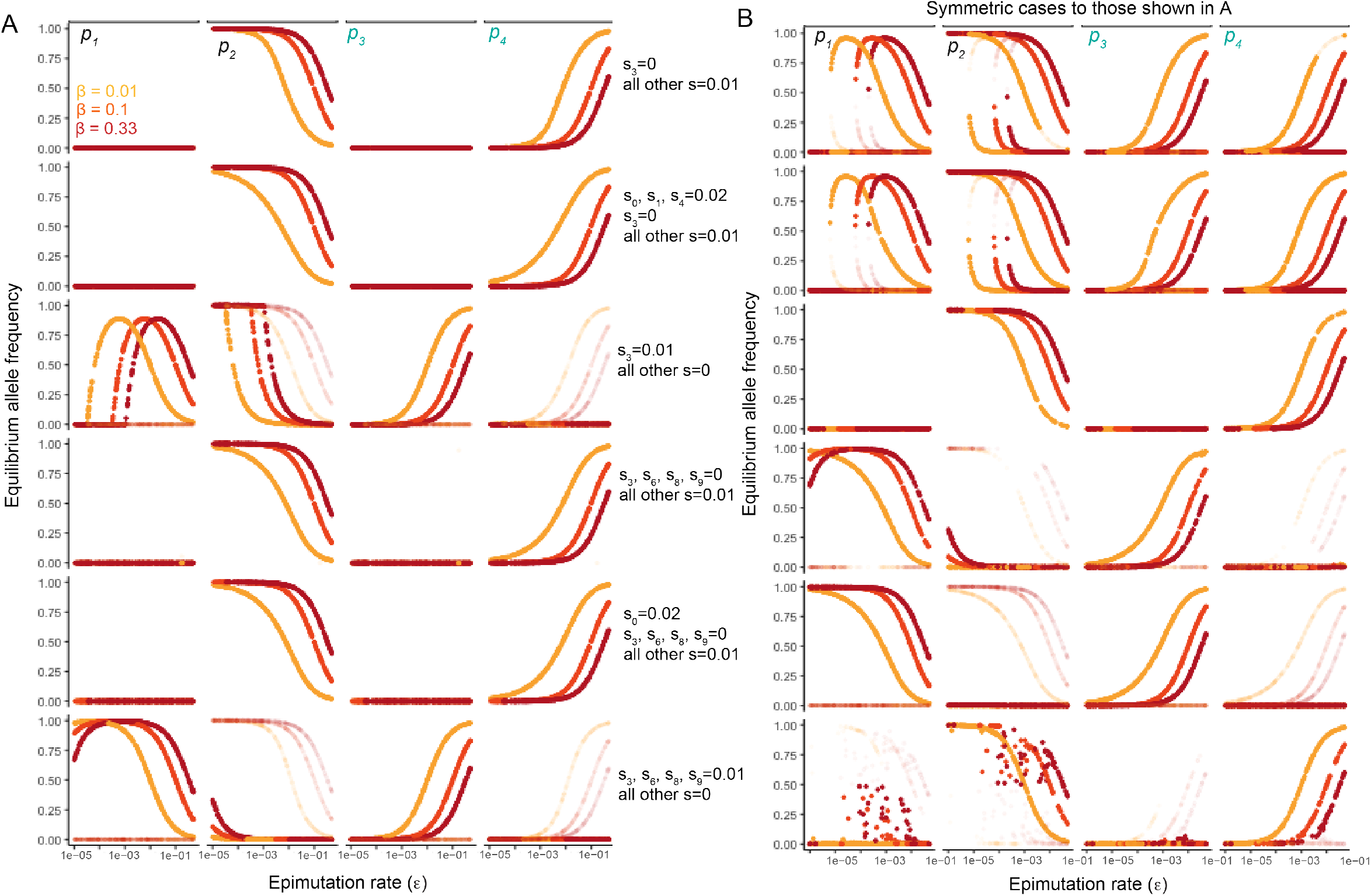
Exact solutions and symmetric cases corresponding to Figure 4. Exact solutions and symmetric cases corresponding to Figure 5. A-B. Equilibrium allele frequencies for *p*_1_ through *p*_4_. Stable equilibria are completely opaque points, while unstable equilibria are faded. A. From top to bottom facet, raw output for the cases shown in Figure 4 and referenced in the corresponding text with corresponding selection coefficients. B. From top to bottom facet, raw output for symmetric values to those shown A. Symmetric means that selection coefficients corresponding to *A*_1_ and *A*_3_ are swapped with *a*_2_ and *a*_4_, respectively.

**Supplementary Figure 2:**
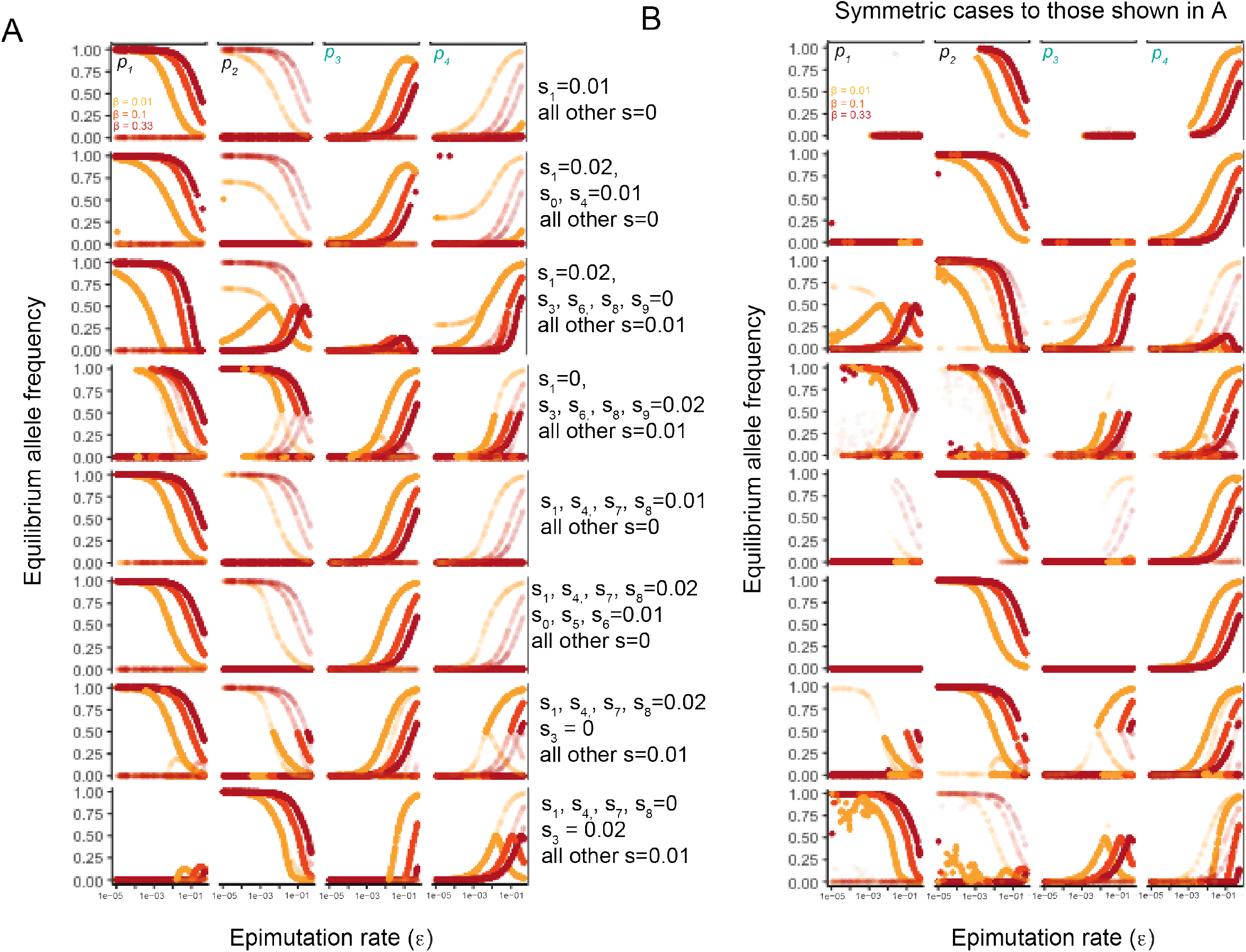
Exact solutions and symmetric cases corresponding to Figures 5 and 6. A-B. Equilibrium allele frequencies for *p*_1_ through *p*_4_. Stable equilibria are points with complete opacity, while unstable equilibria are faded. A. From top to bottom facet, raw output for cases shown in Figures 5 and 6 and referenced in the text, with corresponding selection coefficients. B. From top to bottom facet, raw output for symmetric values to those in A. Symmetric means that selection coefficients corresponding to *A*_1_ and *A*_3_ are swapped with *a*_2_ and *a*_4_, respectively.

## DATA AVAILABILITY STATEMENT

Mathematica scripts for the full parameter space and focal back-epimutation rates, in addition to csv files and R scripts used to generate plots, are available on Github (https://github.com/amykwebster/2022_EpigeneticModels). Upon publication, the final version on Github will be used to generate a unique DOI on Zenodo.

## ACKNOWLEDGEMENTS

We thank members of the Phillips lab and anonymous reviewers for critical feedback. Funding for this work was provided by NIH grant number R35GM131838 to PCP and F32GM146402 to AKW.

